# Loss of ASD-Related Molecule Cntnap2 Affects Colonic Motility in Mice

**DOI:** 10.1101/2023.04.17.537221

**Authors:** Beatriz G. Robinson, Beau A. Oster, Keiramarie Robertson, Julia A. Kaltschmidt

## Abstract

Gastrointestinal (GI) symptoms are highly prevalent among individuals with autism spectrum disorder (ASD), but the molecular link between ASD and GI dysfunction remains poorly understood. The enteric nervous system (ENS) is critical for normal GI motility and has been shown to be altered in mouse models of ASD and other neurological disorders. Contactin-associated protein-like 2 (Cntnap2) is an ASD-related synaptic cell-adhesion molecule important for sensory processing. In this study, we examine the role of Cntnap2 in GI motility by characterizing Cntnap2’s expression in the ENS and assessing GI function in *Cntnap2* mutant mice. We find Cntnap2 expression predominately in enteric sensory neurons. We further assess *in-vivo* and *ex-vivo* GI motility in *Cntnap2* mutants and show altered transit time and colonic motility patterns. The overall organization of the ENS appears undisturbed. Our results suggest that Cntnap2 plays a role in GI function and may provide a molecular link between ASD and GI dysfunction.

## 1 Introduction

Autism spectrum disorder (ASD) is a neurodevelopmental disorder affecting approximately 1 in 36 children in the USA (Maenner et al., 2023). Individuals with ASD often report gastrointestinal (GI) issues, which can lead to irritability and social withdrawal, ultimately affecting quality of life (Chaidez et al., 2014; Restrepo et al., 2020; Valicenti-McDermott et al., 2006). GI issues, including constipation, diarrhea, and abdominal pain (Chaidez et al., 2014; Holingue et al., 2018), have been correlated with sensory over-responsivity in the central and peripheral nervous system in children with ASD (Mazurek et al., 2013). Whether sensory functions of the GI tract are also affected in ASD, and thus potentially contribute to ASD-related GI dysfunction, has not yet been extensively explored.

The enteric nervous system (ENS) is a quasi-autonomous neuronal network that populates the length of the GI tract and can regulate GI function and motility independent of the central nervous system (CNS) (Furness et al., 1994). Enteric neurons and glia cluster into ganglia that reside within the myenteric and submucosal plexuses within the gut wall (Sasselli et al., 2012). GI motility initiates when intrinsic enteric sensory neurons, known as intrinsic primary afferent neurons (IPANs), are activated by chemical or mechanical stimuli. IPANs signal to enteric interneurons that stimulate excitatory or inhibitory motor neurons, resulting in repetitive contractions and propulsive motility (Fung & Berghe, 2020; Furness et al., 1994; Rao & Gershon, 2016). Alterations in ENS activity, organization or gene expression are known to affect digestive function (Rao & Gershon, 2016; Avetisyan et al., 2015). We hypothesize that genes known to be risk factors for ASD are expressed in the ENS and influence enteric neuron activity, and thus could provide a link between ASD and associated GI dysfunction.

ASD-related genes have previously been linked to GI function (Niesler & Rappold, 2021). Mutations in the zebrafish *shank3* gene, which encodes a synaptic scaffolding protein critical for synaptic transmission, result in reduced serotonin-expressing enteroendocrine cells and serotonin-filled ENS boutons, and prolonged GI transit (James et al., 2019). In mice, a global deletion of Nlgn3, an ASD-related synaptic cell adhesion molecule, results in increased colonic diameter and faster colonic migrating motor complexes (Leembruggen et al., 2019).

Here we study Contactin-associated protein-like 2 (Cntnap2; also known as Caspr2), an ASD-related cell-adhesion molecule that aids in the formation and function of the central and peripheral nervous system (Anderson et al., 2012; Gordon et al., 2016; Peñagarikano & Geschwind, 2012; Poliak et al., 1999). *CNTNAP2* gene mutations have been detected in individuals diagnosed with ASD (Peñagarikano & Geschwind, 2012), and *Cntnap2*^*-/-*^ mice show social deficits, communication impairment, and repetitive behaviors, three hallmark characteristics of ASD (Peñagarikano et al., 2011). Additionally, *Cntnap2*^*-/-*^ mice have altered neural circuitry in the somatosensory cortex and exhibit hypersensitivity to mechanical stimuli due to enhanced excitability of primary dorsal root afferents (Dawes et al., 2018; Peñagarikano et al., 2011). In the GI tract, CNTNAP2 has been associated with inflammatory bowel disease, and *Cntnap2*^*-/-*^ mice have increased intestinal permeability (Buniello et al., 2019; Graf et al., 2019). Whether GI motility, which relies on sensing luminal stimuli, is affected in *Cntnap2*^*-/-*^ mice has not been previously investigated.

In this study, we assess Cntnap2 expression in the adult mouse GI tract and ask whether ENS organization and GI motility are altered in *Cntnap2*^*-/-*^ mice. We find that Cntnap2 is predominantly expressed in IPANs, being nearly exclusive to IPANs in the colon. We assess GI motility *in-vivo* and focus on colonic motor function in an *ex-vivo* motility monitor in the absence and presence of an artificial stimulus. We find that lack of Cntnap2 results in altered colonic motility. The overall organization of the ENS appears undisturbed.

## 2 Material and Methods

### 2.1 Animals

All procedures conformed to the National Institutes of Health Guidelines for the Care and Use of Laboratory Animals and were approved by the Stanford University Administrative Panel on Laboratory Animal Care. C57BL/6, B6.129(Cg)-Cntnap2^tm1Pele^/J (Strain #:017482, hereafter *Cntnap2*^*-*^), and B6.129(Cg)-Cntnap2^tm2Pele^/J (Strain #:028635, hereafter *Cntnap2*^*tlacZ*^) mice were purchased from The Jackson Laboratory. Mice were maintained on a 12:12 LD cycle and fed a standard rodent diet, containing 18% Protein and 6% Fat (Envigo Teklad). Food and water were provided ad libitum and mice were group housed with a maximum of five adults per cage. Both male and female 8-12 week-old adult mice were used in this study.

### 2.2 Histology

Mice were euthanized by CO_2_ followed by cervical dislocation. Segments of SI and colon were dissected, flushed with cold PBS, and cut longitudinally along the mesenteric border. Segments were opened flat, placed between sheets of filter paper, and immersed in 4% PFA at 4°C for 90 min. Tissue was rinsed three times in PBS for 10 min and immersed in a 30% sucrose solution overnight at 4°C. Tissue sections were rolled into a “swiss-roll” preparation as described (Williams et al., 2016), embedded in OCT (Tissue-Tek), and frozen until use. 14 μm slices were sectioned using a Leica Cryostat (Leica CM3050 S) and mounted on Superfrost glass slides. Slides were stained with hematoxylin and eosin (H&E). Brightfield images were taken by the Human Pathology/Histology Service Center at Stanford School of Medicine and analyzed for villus height, crypt depth, colonic fold thickness, and circular muscle thickness using Leica ImageScope software. Villus height was measured when full lacteal was visible and crypt depth was measured when both villus/crypt junctions were present in the jejunum. Colonic fold thickness was measured from cross sections of mid and distal colon. 10 measurements were taken per animal. To determine muscle thickness, 20 measurements were taken at random points along the length of the jejunum and distal colon.

### 2.3 Tissue Dissection and Processing

Dissection and tissue processing of the intestines was performed as previously described (Hamnett et al., 2022). Wholemount muscle-myenteric plexus preparations were made by peeling away the muscularis (longitudinal and circular muscle with myenteric plexus). The tissue was stored in PBS with 0.1% sodium azide at 4°C for up to 3 weeks. Jejunum samples were taken from the middle 1/3 length of the SI. The final 1/3 of the colon was considered distal colon.

### 2.4 Immunohistochemistry

Segments of the jejunum (≥1 cm in length) and distal colon (≥0.5 cm in length) were used for immunohistochemistry studies. Staining was performed as previously described (Hamnett et al., 2022), with modifications. For cell body labeling with anti-Cntnap2 antibody, PBT contained 0.01% Triton X-100; for all other labeling, PBT contained 0.1% Triton X-100 (Figure S1C-D). Primary antibodies used included rabbit anti-Cntnap2 (1:1000; Alomone Labs, APZ-005), rabbit anti-ß-galactosidase (1:1000; gift from J. Sanes), human anti-HuC/D (ANNA1) (1:100,000; gift from V. Lennon), goat anti-Sox10 (1:2,000; R&D Systems, AF2864) and fluorophore-conjugated secondary antibodies (Jackson Labs and Molecular Probes).

### 2.5 RNAscope *In Situ* Hybridization with Protein Co-detection

Tissue was dissected and prepared for fixation as outlined in *Tissue Dissection and Processing*. Flat segments of the jejunum and distal colon were fixed overnight in 4% PFA. Segments were rinsed with PBS, and wholemount muscle-myenteric plexus preparations were made by peeling away the muscularis. RNAscope *in situ* with protein co-detection was performed using Advanced Cell Diagnostics (ACD) RNAscope Multiplex Fluorescent Reagent Kit v2 (Cat# 323100) and ACD RNA-protein Co-detection ancillary kit (Cat# 323180) as described in (Guyer et al., 2023). The following RNAscope probes were used: Mm-Nmu-C1 (Cat# 446831) and Mm-Cntnap2-C1 (Cat# 449381).

### 2.6 Neuron Quantification

Images were acquired on a Leica SP8 confocal microscope using 20x and 63x oil objectives. All images were adjusted for brightness and contrast using ImageJ/FIJI. For Cntnap2 quantification, three 20x ROIs (1000 μm x 1000 μm) per mouse were randomly selected in both the jejunum and distal colon. HuC/D^+^ and Cntnap2^+^ neurons were counted manually using the cell counter tool FIJI (Schindelin et al., 2012). For each region, neurons per ROI were averaged per animal.

For quantification of Cntnap2 co-expression with *Nmu* transcript, five images (138 μm x 138 μm) were taken at 63x magnification per region per mouse. Regions of interest (ROIs) were created around every HuC/D^+^ neuron for each image and manually scored as either positive or negative for Cntnap2 or *Nmu* transcript. Neurons with ≥20 *Nmu* or *Cntnap2* fluorescent transcript dots were considered positive. For each region, the average percentage of co-expression was calculated per mouse.

Quantification of ganglia was performed using COUNTEN (Kobayashi et al., 2021), with *σ* = 4.5. For each region, the average of three maximum projection images (1000 μm x 1000 μm) were analyzed per mouse.

### 2.7 Functional Behavior

#### 2.7.1 Whole GI transit time

Whole GI transit assay was performed as previously described (Spear et al., 2018). In brief, mice were gavaged with a carmine red-methylcellulose mixture and observed until a red pellet was expelled.

#### 2.7.2 Gastric Emptying and SI Transit

Gastric emptying and SI transit were determined as previously described (De Lisle, 2007; Spear et al., 2018). In brief, mice were fasted for 12 hours and water was removed 3 hours before the start of the assay. Mice were gavaged with a 2% methylcellulose mixture containing 2.5 mg/mL Rhodamine B Dextran (Invitrogen, D1841, MW: 70,000). 15 minutes after gavage, mice were euthanized with CO_2_ and the stomach and SI were removed. The SI was divided into 10 equal segments that were homogenized in saline. The fluorescence in the stomach and each SI segment was measured. The percentage of gastric emptying and the geometric center were determined as previously described (De Lisle, 2007).

#### 2.7.3 Bead Expulsion Assay

Bead expulsion assay was performed as previously described (Spear et al., 2018). In brief, mice were lightly anesthetized by isoflurane and a 2 mm glass bead was inserted 2 cm into the colon through the anus using a gavage needle. Expulsion time was determined as the time from bead insertion to when the bead was fully expelled.

#### 2.7.4 Fecal Water Content & Pellet Length

Fecal water content was assessed as previously described (Spear et al., 2018) with modifications to allow for measurement of pellet lengths. Mice were housed individually for 1 hour during which all fecal pellets were collected immediately after expulsion, photographed, and stored in a pre-weighed tube (1 tube/mouse). After 1 hour of collection, tubes were weighed again, incubated for 48 hours at 50°C, and weighed a final time to determine the percentage of water content. Pellet length was measured using FIJI (Swaminathan et al., 2019).

### 2.8 *Ex-vivo* Colonic Motility Assay

*Ex-vivo* motility monitor assay was adapted from (Hennig et al., 1999; Spear et al., 2018; Swaminathan et al., 2016). Colons with cecum attached were removed and placed in warmed Kreb’s solution. The mesentery was cut away, and colons were placed in an organ bath, pinned down at the cecum and distal colon end with care to not impede expulsion of contents. The organ bath was kept at 37°C and filled with circulating warmed Kreb’s solution (NaCl, 120.9 mM; KCl, 5.9 mM; NaHCO_3_, 25.0 mM; Monobasic NaH_2_PO_4_, 1.2 mM; CaCl_2_, 3.3 mM; MgCl_2_•6H_2_0, 1.2 mM; D-Glucose, 11.1 mM) saturated with carbogen (95% O_2_ and 5% CO_2_). Colons were allowed to acclimate for 10 minutes in the bath. Colonic motility was recorded *ex-vivo* using a high-resolution monochromatic firewire industrial camera (The Imaging Source, DMK41AF02) mounted directly above the organ bath as previously described (Spear et al., 2018; Swaminathan et al., 2016).

#### 2.8.1 Motility Monitor – Natural Colonic Behavior

After a 10-minute acclimation period and additional 20-minutes to allow for clearing of natural fecal pellets, motility of the empty colon was recorded for a 10-minute period. Recorded videos were converted to spatiotemporal maps (STMs) using Scribble 2.0 and Matlab (2012a) plugin Analyse 2.0 (Swaminathan et al., 2016) and annotated to determine characteristics of CMCs, which we considered neurogenic repetitive contractions (Corsetti et al., 2019). Intervals between CMCs were measured from start of one contraction to the start of the next contraction (Fida et al., 2000).

#### 2.8.2 Motility Monitor – Artificial Pellet Assay

Dissection was performed as described in “*Ex-vivo* Colonic Motility Assay”, with cecum removed. Artificial pellet assay was adapted from (Costa et al., 2021). The colon was flushed of endogenous fecal matter using warmed Kreb’s solution. After 10 min of acclimation, a lubricated (KY jelly) 2 mm 3D-printed pellet was inserted through the proximal colon and gently pushed to the proximal-mid colon junction using a blunt-ended gavage needle. Colonic activity was recorded until the pellet was fully expelled from the distal end of the colon. After at least three successful trials in which the artificial pellet traveled through the colon independently and was fully expelled, 10 additional minutes of empty colonic activity were recorded to ensure normal function. Time to expulsion was determined and the pellet’s path was traced using FIJI plug-in TrackMate (v7.6.1), from which pellet velocity and max speed were determined (Ershov et al., 2021; Tinevez et al., 2017). STMs were generated as described above.

### 2.9 Statistical Analysis

Statistical analyses were performed using GraphPad Prism (Version 9.4.1) with a 95% confidence limit (p < 0.05). Data are presented as mean ± SEM and checked for normal distribution. Unless otherwise noted, an unpaired t-test was used for comparison between two groups. For comparison between more than two groups, one-way or two-way analysis of variance (ANOVA) was used with Tukey’s multiple comparisons test. To ensure sufficient animals were used for the studies, we performed power analyses based on early pilot data using a p-value (alpha) of 0.05 and a power (beta) of 0.8. Experimenter and analyzer were blinded to the genotype when feasible and appropriate. “n” refers to the number of animals tested, unless otherwise stated.

## 3 Results

To define the distribution of Cntnap2 in the mouse intestines, we used an antibody against Cntnap2, which we validated using *Cntnap2* transcript co-expression (Figure 1A) and *Cntnap2*^*-/-*^ mice (Figure 1B) (Poliak et al., 2003). As we were interested in querying the role of Cntnap2 in GI motility, we focused our analysis on the myenteric plexus, which harbors the intrinsic neuronal circuitry required for motility (Spencer & Hu, 2020). We examined *Cntnap2*^*tlacz/+*^ mice (Gordon et al., 2016), and found β-gal-expressing neurons and projections throughout the SI and colon (Figure 1C and data not shown). Given that the expression of neurotransmitters and neuromodulators can differ between intestinal regions (Hamnett et al., 2022), we assessed Cntnap2 expression in distinct regions of the SI (duodenum, jejunum, and ileum) and colon (proximal, mid, and distal) (Figure 1D). Cntnap2 is expressed in 10-25% of HuC/D^+^ enteric neurons, depending on the region analyzed (Figure 1G). We further observed Cntnap2 expression in a small subset of Sox10^+^ progenitor/glial cells (Morarach et al., 2021) (Figure S1A) and in some 5-HT^+^ intestinal epithelial cells, suggesting that Cntnap2 is present in a subset of enterochromaffin cells along the epithelial layer (Figure S1B).

**Figure 1.**
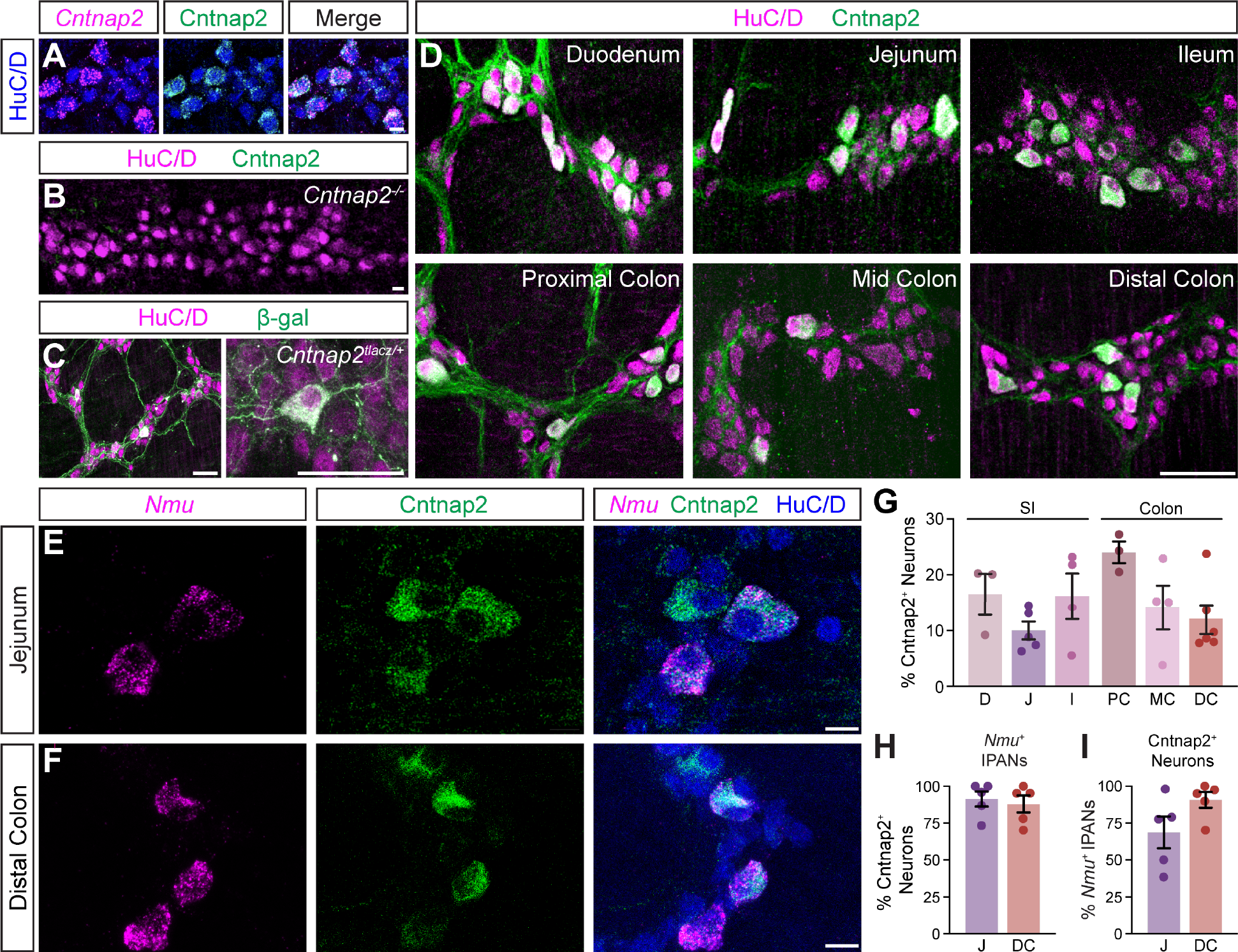
Cntnap2 expression in enteric sensory neurons. **(A)** Cntnap2 (green) colocalizes with *Cntnap2* transcript (magenta) in adult jejunum myenteric plexus. Enteric neurons labeled with HuC/D (blue). **(B)** Cntnap2 (green) is absent from HuC/D^+^ (magenta) enteric neurons in *Cntnap2*^*-/-*^ myenteric plexus of adult jejunum. **(C)** β-gal (green) expression in HuC/D^+^ (magenta) neurons of adult *Cntnap2*^*tlacZ/+*^ jejunum. **(D)** Cntnap2 (green) is expressed in a subset of HuC/D^+^ (magenta) neurons throughout the small intestine and colon. **(E, F)** A subset of *Nmu*^*+*^ (magenta) sensory neurons expresses Cntnap2 (green) in the jejunum (E) and distal colon (F). **(G)** Quantification of Cntnap2^+^ HuC/D^+^ neurons in the small intestine (D: 16.4 ± 3.6% [n = 3]; J: 10.0 ± 1.6% [n = 5]; I: 16.1 ± 4.0% [n = 4]) and colon (PC: 23.9 ± 1.9% [n = 3]; MC: 14.2 ± 3.9% [n = 4]; DC: 11.9 ± 2.5% [n = 6]). **(H)** The majority of *Nmu*^+^ IPANs express Cntnap2 in the jejunum (91.2 ± 5.1% [n = 5]) and distal colon (87.8 ± 5.7% [n = 5]). **(I)** The majority of Cntnap2^+^ neurons express *Nmu* in the jejunum (68.6 ± 10.7% [n = 5]) and distal colon (90.5 ± 5.3% [n = 5]). Scale bars, (A, B, E, F) 10 μm, (C, D) 50 μm. D: Duodenum, J: Jejunum, I: Ileum, PC: Proximal colon, MC: Mid colon, DC: Distal colon.

We next asked whether Cntnap2 expression in the myenteric ENS was confined to a particular neuronal subtype. We focused this and all future analyses on the distal region of the colon due to its association with the propulsion of formed fecal pellets, and for comparison, chose the jejunum as a representative region within the SI. scRNA-sequencing studies of the mouse ENS have reported high Cntnap2 expression in putative sensory neuron populations in both the SI and colon (Drokhlyansky et al., 2020; Morarach et al., 2021; Zeisel et al., 2018). IPANs make up approximately 26% of enteric neurons in the SI and have Dogiel Type II morphology, based on their large and smooth cell bodies and two or more long axons (Qu et al., 2008). We observed that the majority of Cntnap2^+^ neurons were large in shape with smooth cell bodies (Figure S1A). We further assessed Cntnap2 expression in IPANs, using *Nmu* transcript as a sensory neuron marker (Morarach et al., 2021) (Figure 1E, F). Over 80% of *Nmu*^*+*^ neurons in both SI and colon co-expressed Cntnap2 (Figure 1H) and over half of Cntnap2^+^ neurons in the SI and over 80% of Cntnap2^+^ neurons in the colon colocalized with *Nmu* (Figure 1I). Taken together, these results suggest that Cntnap2 has a subtype and region-specific expression profile, and that the majority of Cntnap2^+^ myenteric plexus neurons in the colon are putative sensory neurons.

We next asked whether the absence of Cntnap2 affects GI morphology and function. *Cntnap2*^*-/-*^ mice survive (Poliak et al., 2003), have normal body weight and SI and colon length (Figure 2A-C). We found no changes in SI villi height, crypt depth, and circular muscle thickness (Figure 2D, E, G, H). Also, colonic fold depth and circular muscle thickness was the same in *Cntnap2*^*-/-*^ as compared to *Cntnap2*^*WT*^ mice (Figure 2F-H). To assess whole GI transit time, we measured the length of time needed for a carmine red mixture gavaged into the stomach to be expelled as a red fecal pellet (Spear et al., 2018). We observed a 23% increase in whole GI transit time when comparing *Cntnap2*^*-/-*^ to *Cntnap2*^*WT*^ mice (Figure 2I). We found no changes in fecal water content and pellet length (Figure 2J, K). Thus, whole gut transit is prolonged in the absence of Cntnap2.

**Figure 2.**
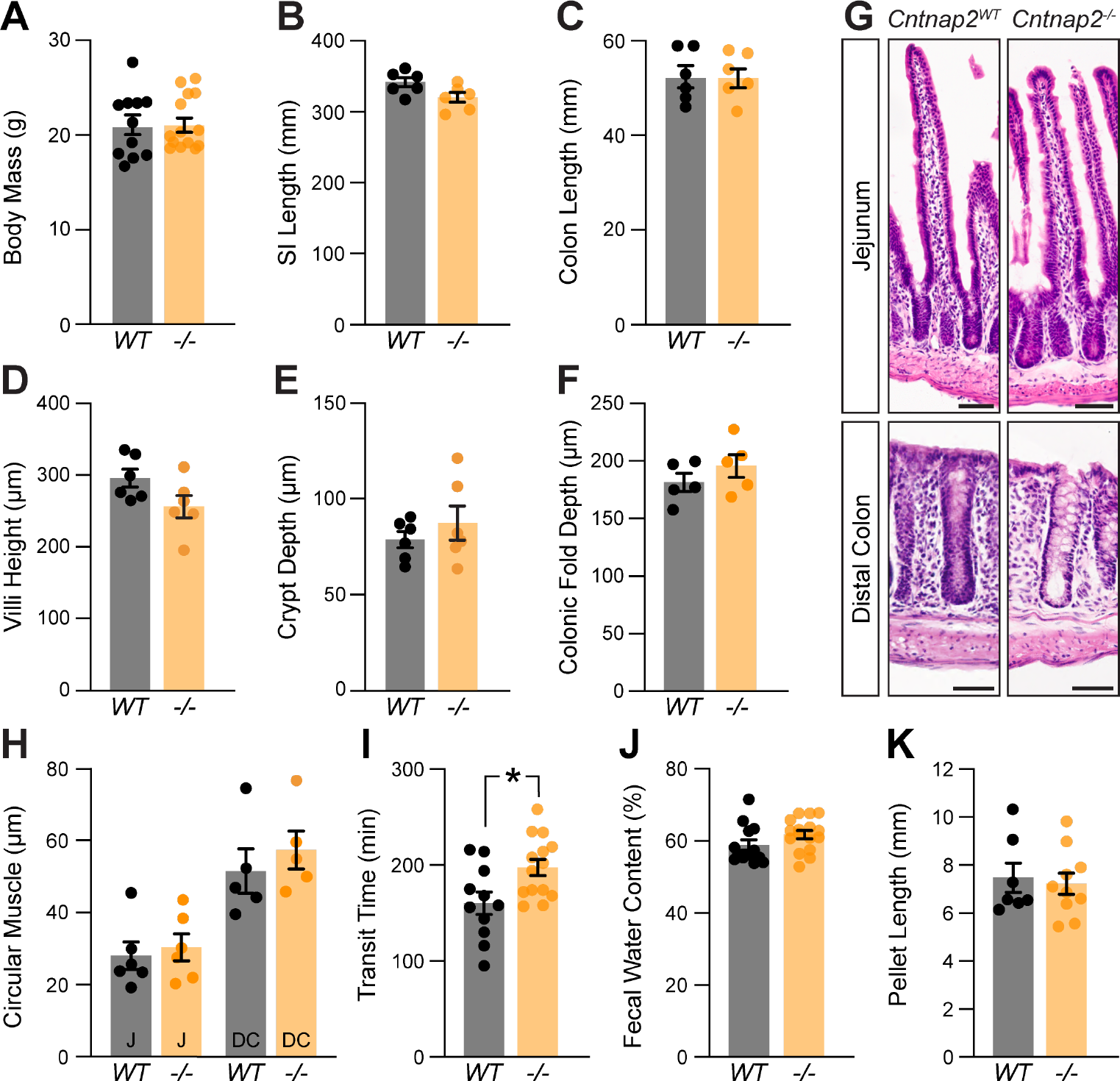
A role for Cntnap2 in whole GI transit. **(A)** Body mass is the same in *Cntnap2*^*WT*^ and *Cntnap2*^-/-^ mice (WT: 21.1 ± 1.0 g [n = 11]; -/-: 21.3 ± 0.8 g [n = 14]). Unpaired t-test, *P =* 0.89. **(B)** Length of small intestine is the same in *Cntnap2*^*WT*^ and *Cntnap2*^-/-^ mice (WT: 341.5 ± 6.5 mm [n = 6]; -/-: 319.8 ± 7.0 mm [n = 6]). Unpaired t-test, *P =* 0.05. **(C)** Length of colon is the same in *Cntnap2*^*WT*^ and *Cntnap2*^-/-^ mice (WT: 52.5 ± 2.3 mm [n = 6]; -/-: 52.4 ± 1.9 mm [n = 6]). Unpaired t-test, *P =* 0.98. **(D)** Villi height is the same in *Cntnap2*^*WT*^ (294.8 ± 12.5 μm [n = 6]) and *Cntnap2*^-/-^ (256.1 ± 15.6 μm [n = 6]) jejunum. Unpaired t-test, *P =* 0.08. **(E)** Crypt depth is the same in *Cntnap2*^*WT*^ (78.6 ± 4.2 μm [n = 6]) and *Cntnap2*^-/-^ (87.6 ± 8.8 μm [n = 6]) mice. Unpaired t-test, *P =* 0.38. **(F)** Depth of colonic folds is the same in *Cntnap2*^*WT*^ (181.4 ± 7.8 μm [n = 5]) and *Cntnap2*^-/-^ (195.9 ± 10.3 μm [n = 5]) mice. Unpaired t-test, *P =* 0.29. **(G)** H&E stained cross sections of jejunum and distal colon from *Cntnap2*^*W*T^ and *Cntnap2*^*-/-*^ mice. **(H)** Circular muscle thickness is the same in *Cntnap2*^*WT*^ and *Cntnap2*^-/-^ jejunum (WT: 27.9 ± 3.8 μm [n = 6]; -/-: 30.2 ± 3.7 μm [n = 5]) and distal colon (WT: 51.5 ± 6.1 μm [n = 5]; -/-: 57.4 ± 5.3 μm [n = 5]). Two-way ANOVA: genotype, F(1,18) = 0.77, *P* = 0.39; region, F(1,18)= 29.12, *P <* 0.001; interaction, F(1,18)= 0.14, *P =* 0.71. **(I)** Whole GI transit time is increased in *Cntnap2*^-/-^ (197.6 ± 8.5 min, [n = 14]) compared to *Cntnap2*^*WT*^ (160.5 ± 11.6 min [n = 11]) mice. Unpaired t-test, *P =* 0.01. **(J)** Fecal water content is the same in 9 week old *Cntnap2*^*WT*^ (58.8 ± 1.5% [n = 13]) and *Cntnap2*^-/-^(61.8 ± 1.1% [n = 16]) mice. Unpaired t-test, *P =* 0.12. **(K)** Fecal pellet length is the same in 9-week-old *Cntnap2*^*WT*^ (7.5 ± 0.6 mm [n = 7]) and *Cntnap2*^-/-^ (7.2 ± 0.4 mm [n = 10]) mice. Unpaired t-test, *P =* 0.73. All mice were 11 weeks old unless stated otherwise. Tukey’s multiple comparison test: * p < 0.05, ** p < 0.01, *** p < 0.001. Scale bar, 50 μm. WT: *Cntnap2*^*WT*^; -/-: *Cntnap2*^*-/-*^; SI: Small Intestine; J: Jejunum; DC: Distal Colon.

Whole gut transit provides information about stomach, small intestine and colon transit combined. To specifically focus on the colon, we next recorded and analyzed colonic motility using an *ex-vivo* motility monitor (Swaminathan et al., 2016). In this setup, the colon is isolated from extrinsic innervation and allows us to assess loss of GI tract intrinsic Cntnap2. We generated spatiotemporal maps (STMs) of the empty colon (Figure 3A) and observed that CMCs were 31% shorter-lasting in *Cntnap2*^*-/-*^ compared to *Cntnap2*^*WT*^ mice (Figure 3F, G). CMC intervals, number, velocity, and length remained unchanged (Figure 3B-E, G). Thus, repetitive contractions are shortened in isolated empty *Cntnap2* mutant colons.

**Figure 3.**
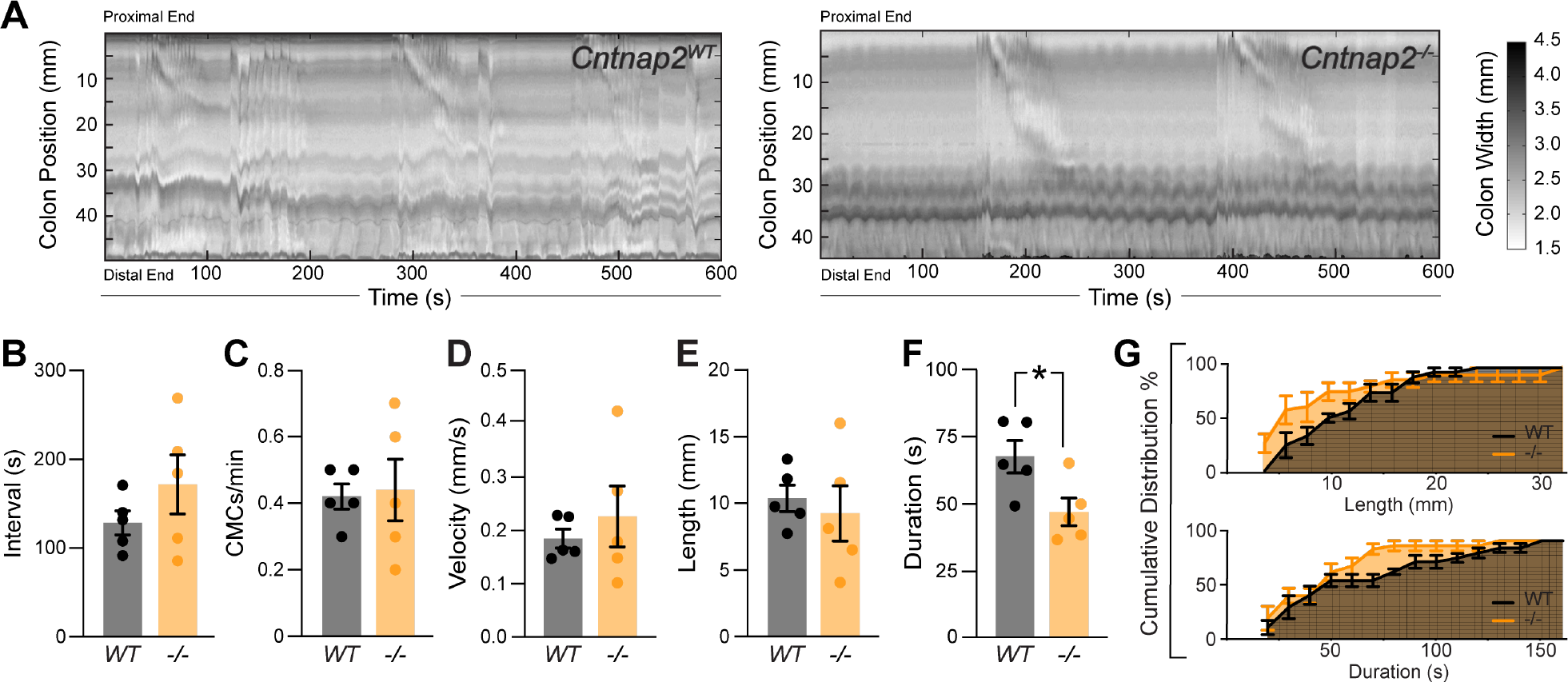
Altered *ex-vivo* motility in empty *Cntnap2*^*-/-*^ colons. **(A)** Representative spatiotemporal maps of 10 minute video recordings from *Cntnap2*^*WT*^ and *Cntnap2*^*-/-*^empty colons. Gray scale indicates colonic diameter. **(B)** Intervals between CMC onsets are unchanged in *Cntnap2*^-/-^ (171.7 ± 33.2 s [n = 5]) compared to *Cntnap2*^*WT*^ (128.8 ± 13.7 s [n = 5]) mice. Unpaired t-test, *P =* 0.27. **(C)** Number of CMCs per minute are the same in *Cntnap2*^*WT*^ (0.42 ± 0.04 [n = 5]) and *Cntnap2*^-/-^ (0.44 ± 0.09 [n = 5]) mice. Unpaired t-test, *P =* 0.84. **(D)** CMC velocity is the same in *Cntnap2*^*WT*^ (0.18 ± 0.02 mm/s [n = 5]) and *Cntnap2*^-/-^ (0.23 ± 0.06 mm/s [n = 5]) mice. Unpaired t-test, *P =* 0.49. **(E)** Length of CMCs is the same in *Cntnap2*^*WT*^ (10.3 ± 1.0 mm, [n = 5]) and *Cntnap2*^-/-^ (9.3 ± 2.1 mm [n = 5]) mice. Unpaired t-test, *P =* 0.66. **(F)** CMCs are shorter-lasting in *Cntnap2*^-/-^ (47.6 ± 5.2 s [n = 5]) compared to *Cntnap2*^*WT*^ (68.5 ± 6.0 [n = 5]) mice. Unpaired t-test, *P =* 0.03. **(G)** Cumulative distributions for CMC length (mm) and duration (s) in *Cntnap2*^*WT*^ and *Cntnap2*^*-/-*^ mice.

Given that IPANs are sensitive to stretch (Furness et al., 2004), we next assessed *ex-vivo* colonic motility of *Cntnap2*^*-/-*^ mice in response to a stimulus. We inserted a natural-shaped 3D-printed artificial fecal pellet through the proximal colon and recorded colonic behavior until complete pellet expulsion (Costa et al., 2021). The artificial pellet served as a normalized stimulus that was able to travel the entire length of the mid and distal colon (Figure 4A, B). The time to pellet expulsion was shortened by 51% in *Cntnap2*^*-/-*^ mice compared to *Cntnap2*^*WT*^ littermate controls (Figure 4C). Using TrackMate (v7.6.1) (Ershov et al., 2021; Tinevez et al., 2017) to create a trace of the pellet’s movement (Figure 4A’), we observed a 42% reduction in the number of pellet movements in *Cntnap2*^*-/-*^ compared to *Cntnap2*^*WT*^ mice (Figure 4D), which was also visible when plotting mean pellet speed along the length of the colon (Figure 4E). The mean pellet speed per trial trended higher in *Cntnap2*^*-/-*^ mice (Figure 4F), but was not statistically significant. Maximum pellet speed was unchanged (Figure 4G). Thus, artificial fecal pellets move more continuously, and colonic transit is accelerated in isolated *Cntnap2* mutant colons.

**Figure 4.**
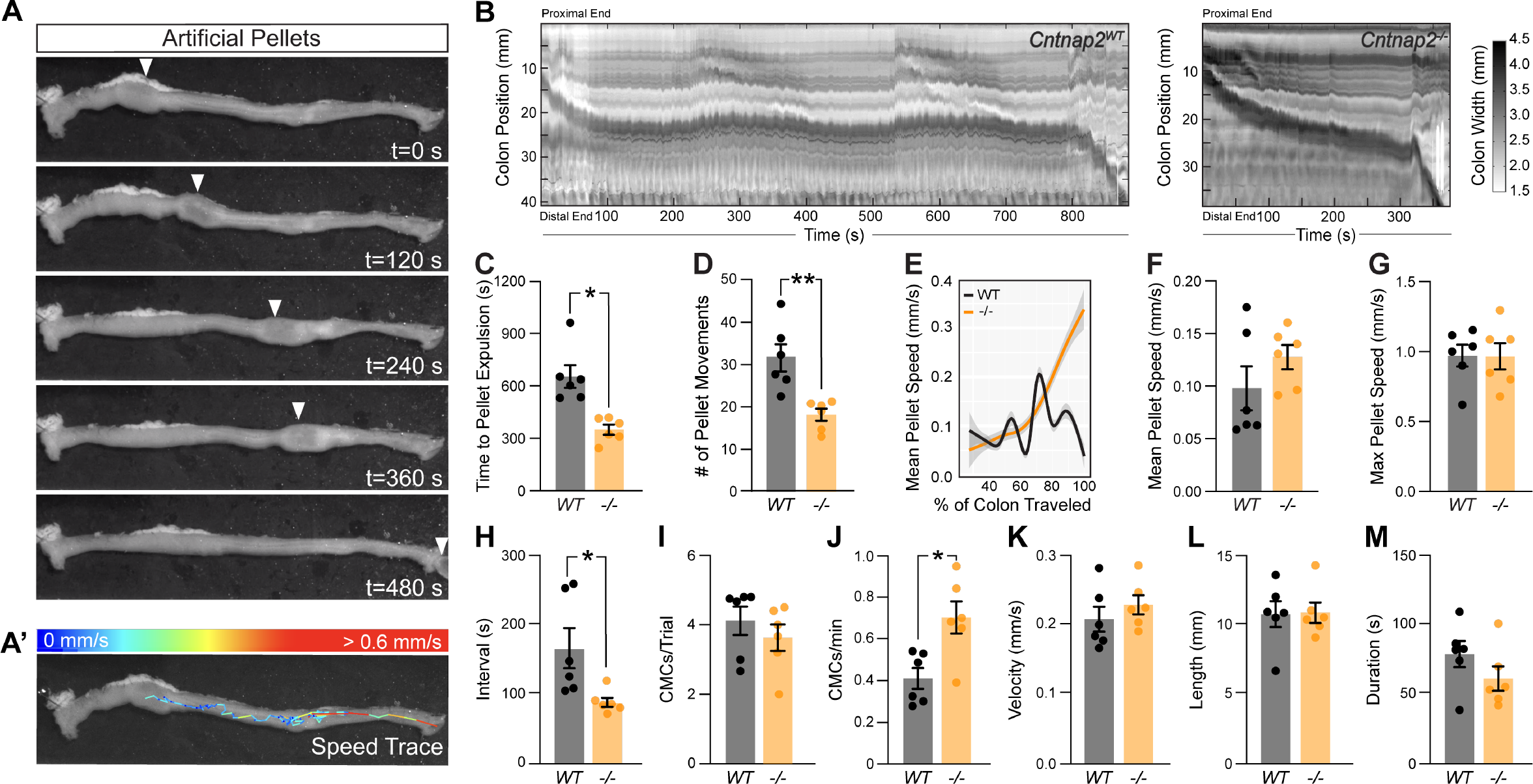
Accelerated *ex-vivo* pellet expulsion in *Cntnap2*^-/-^ colons. **(A)** Time series images of artificial pellet traversing the colon over time. White arrowheads indicate center of artificial pellet. **(A’)** Trace of pellet path with speed of pellet represented by color scale. **(B)** Representative spatiotemporal maps of full-length artificial pellet trials from *Cntnap2*^*WT*^ and *Cntnap2*^*-/-*^ mice. Gray scale indicates colonic diameter. **(C)** Time to pellet expulsion is shorter in *Cntnap2*^*-/-*^ mice (350.2 ± 28.2 s [n = 6]) compared to *Cntnap2*^*WT*^ (657.9 ± 65.0 s [n = 6]) mice. Unpaired t-test, *P =* 0.002. **(D)** Number of pellet movement intervals is significantly reduced in *Cntnap2*^*-/-*^ (18.2 ± 1.4 [n = 6]) when compared to *Cntnap2*^*WT*^ (31.5 ± 3.2 [n = 6]) mice. Unpaired t-test, *P =* 0.003. **(E)** Local regression (LOESS) of mean pellet speed for each genotype as a function of the % of colon length traveled. 95% confidence interval shown in gray. **(F)** Mean speed of artificial pellet per trial is the same in *Cntnap2*^*WT*^ (0.10 ± 0.02 mm/s [n = 6]) and *Cntnap2*^*-/-*^ (0.13 ± 0.01 mm/s [n = 6]) mice. Unpaired t-test, *P =* 0.24. **(G)** Max speed of pellet is the same in *Cntnap2*^*WT*^ (0.97 ± 0.08 mm/s [n = 6]) and *Cntnap2*^*-/-*^ (0.97 ± 0.09 mm/s [n = 6]) mice. Unpaired t-test, *P =* 0.96. **(H)** Intervals between CMCs are significantly reduced in *Cntnap2*^-/-^ (87.1 ± 6.6 s [n = 6]) compared to *Cntnap2*^*WT*^ (165.3 ± 29.0 s [n = 6]) mice. Unpaired t-test, *P =* 0.03. **(I)** Number of CMCs during trial period are similar between *Cntnap2*^*WT*^ (4.1 ± 0.4 [n = 6]) and *Cntnap2*^-/-^(3.6 ± 0.4 min [n = 6]) mice. Unpaired t-test, *P =* 0.4. **(J)** Number of CMCs per minute are increased in *Cntnap2*^-/-^ (0.71 ± 0.08 [n = 6]) compared to *Cntnap2*^*WT*^ (0.41 ± 0.05 [n = 6]) mice. Unpaired t-test, *P =* 0.0099. **(K)** Velocity of CMCs is the same in *Cntnap2*^*WT*^ (0.21 ± 0.02 mm/s [n = 6]) and *Cntnap2*^-/-^ (0.23 ± 0.01 mm/s [n = 6]) mice. Unpaired t-test, *P =* 0.38. **(L)** Length of CMCs is the same in *Cntnap2*^*WT*^ (10.8 ± 0.9 mm [n = 6]) and *Cntnap2*^-/-^ (10.9 ± 0.7 mm [n = 6]) mice. Unpaired t-test, *P =* 0.93. **(M)** Duration of CMCs is the same in *Cntnap2*^*WT*^ (79.1 ± 9.4 s [n = 6]) and *Cntnap2*^-/-^ (61.5 ± 8.8 s [n = 6]) mice. Unpaired t-test, *P =* 0.2. Tukey’s multiple comparison test: * p < 0.05, ** p < 0.01, *** p < 0.001. WT: *Cntnap2*^*WT*^; -/-: *Cntnap2*^*-/-*^.

We next performed STM analysis of these *ex-vivo* colonic motility data in the presence of a stimulus. The interval between CMC onsets was shortened by 47% in *Cntnap2*^*-/-*^ compared to *Cntnap2*^*WT*^ mice (Figure 4H), while the number of CMCs per trial remained the same (Figure 4I). As a result, CMC frequency was 73% increased in *Cntnap2*^*-/-*^ mice (Figure 4J). CMC velocity, length, and duration remained the same (Figure 4K-M). These findings suggest that in the presence of a luminal stimulus, CMC frequency is increased.

Given the predominant expression of Cntnap2 in IPANs, we next asked whether lack of Cntnap2 impacts IPANs. The total number and distribution of HuC/D^+^ myenteric neurons in the distal colon were unchanged in *Cntnap2*^*-/-*^ mice (Figure 5A-E). Also, the number and organization of *Nmu*^*+*^ neurons were unchanged (Figure 5F-G). Thus, the number of myenteric plexus-resident sensory neurons is unchanged.

**Figure 5.**
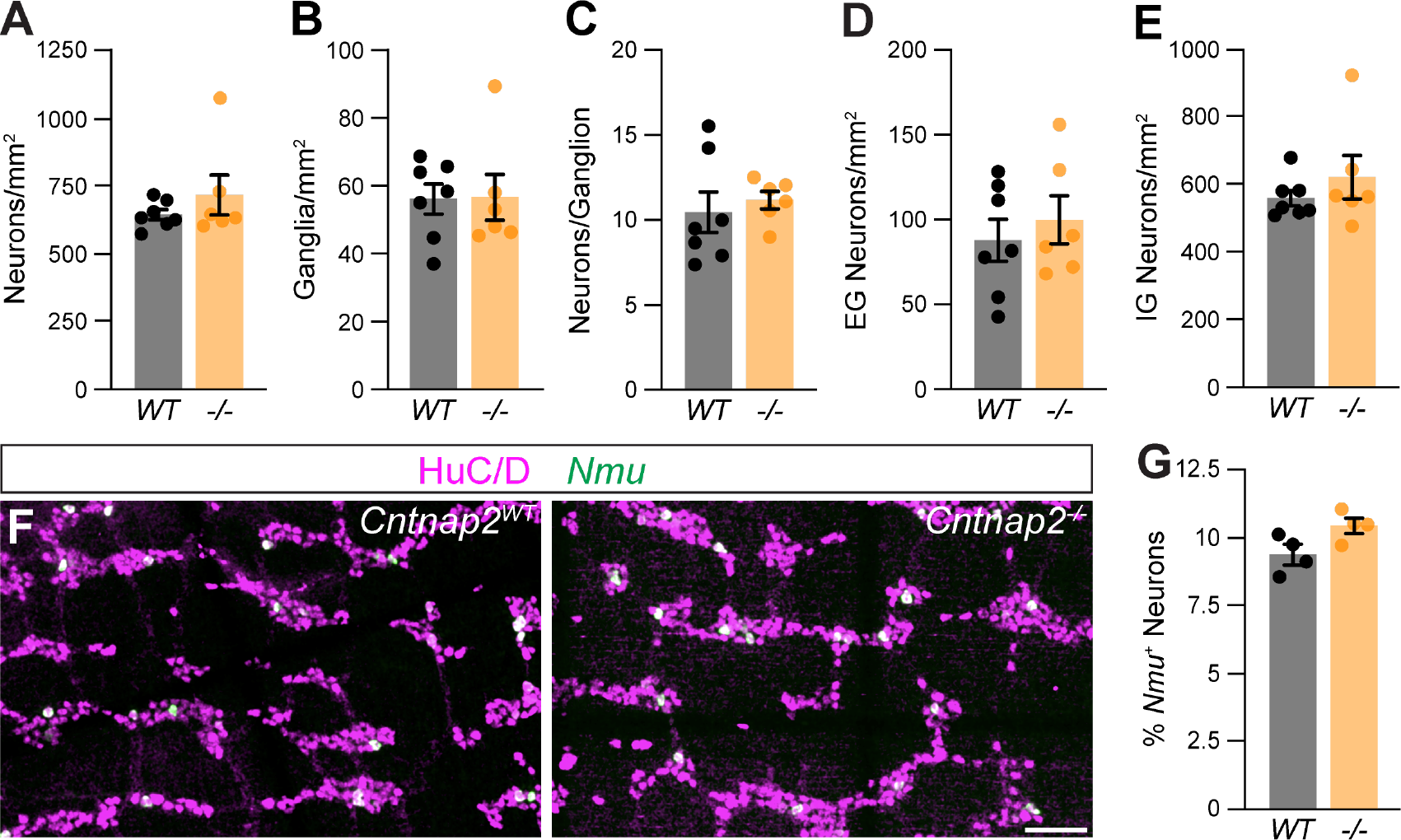
Normal ENS organization in *Cntnap2* mutant distal colon. **(A)** Number of HuC/D^+^ neurons is the same in *Cntnap2*^*WT*^ (647.7 ± 18.8 [n = 7]) and *Cntnap2*^*-/-*^ (720.9 ± 73.6 [n = 6]) mice. Unpaired t-test, *P =* 0.32. **(B)** Number of enteric ganglia is the same in *Cntnap2*^*-/-*^ (56.7 ± 6.8 [n = 6]) compared to *Cntnap2*^*WT*^ (56.2 ± 4.4 [n = 7]) mice. Unpaired t-test, *P =* 0.95. **(C)** Number of neurons per ganglion is unchanged in *Cntnap2*^*-/-*^ (11.2 ± 0.5 [n = 6]) compared to *Cntnap2*^*WT*^ (10.5 ± 1.2 [n = 7]) mice. Unpaired t-test, *P =* 0.62. **(D, E)** Number of extra-(D) and intra-ganglionic (E) neurons are similar in *Cntnap2*^*WT*^ (extra: 88.7 ± 12.6; intra: 559.0 ± 22.7 [n = 7]) and *Cntnap2*^*-/-*^ mice (extra: 100.8 ± 14.4; intra: 620.1 ± 23.8 [n = 7]). Unpaired t-test, *P* (extra, intra) *=* 0.54, 0.36. *Nmu* (green) is expressed in a subset of HuC/D^+^ (magenta) neurons in *Cntnap2*^*WT*^ and *Cntnap2*^*-/-*^distal colon. Percent of *Nmu*^+^ HuC/D^+^ neurons is unchanged in *Cntnap2*^*-/-*^ (10.5 ± 0.3 [n = 4]) compared to *Cntnap2*^*WT*^ (9.4 ± 0.3 [n = 4]) mice. Unpaired t-test, *P =* 0.05. Scale bar, 50 μm.

## 4 Discussion

GI dysfunction is a prevalent symptom in individuals with ASD (Holingue et al., 2018). In this study, we aimed to determine whether the ASD-related gene, *Cntnap2*, plays a role in mouse GI function by characterizing Cntnap2’s expression in the intestines and assessing colonic function and ENS organization in *Cntnap2*^*-/-*^ mice. Our findings reveal that Cntnap2 is expressed in colonic sensory neurons, and a subset of progenitor/glial cells and intestinal epithelial cells. Whole gut transit is slowed in *Cntnap2*^*-/-*^ mice and repetitive contractions are shortened in isolated empty *Cntnap2* mutant colons. In the presence of a luminal stimulus, CMC frequency is increased and colonic transit is accelerated in isolated *Cntnap2* mutant colons. The overall organization of the ENS appears unchanged.

Sensory over-responsivity has been correlated with the presence of GI issues in children diagnosed with ASD (Mazurek et al., 2013) and Cntnap2 has been linked to sensory processing deficits and hypersensitivity in the mouse CNS and PNS (Dawes et al., 2018; Fernández et al., 2021; Peñagarikano et al., 2011). Our finding that Cntnap2 is expressed in the majority of *Nmu*^*+*^ IPANs is consistent with scRNA-seq studies reporting a high expression of Cntnap2 in putative enteric sensory neuron classes (Drokhlyansky et al., 2020; Morarach et al., 2021; Zeisel et al., 2018). IPANs are thought to be critical for initiating propulsive CMCs and downstream motility patterns (Feng et al., 2022; Fung et al., 2021; Nestor-Kalinoski et al., 2021; Spencer et al., 2018) and the observed changes to CMCs in *ex-vivo Cntnap2*^*-/-*^ colon preparations suggest a role for Cntnap2 in enteric sensory function.

One caveat of this study is that the germline Cntnap2 deletion model deletes Cntnap2 not only in intrinsic ENS cells, but also from a small number of progenitor/glial cells and EECs that might contribute to the observed phenotypes (Rao et al., 2017; Servin-Vences et al., 2022). Further, in the SI, Cntnap2 might be additionally expressed in other ENS subsets, such as cholinergic interneurons (Morarach et al., 2021; Zeisel et al., 2018).

Recent studies have provided growing evidence for the role of ASD-related genes in GI function. Other ASD mouse models that have been used to investigate ENS organization include *Slc6a4*^*-/-*^ (SERTKO) mice, SERT Ala56 mice (common SERT variant), *Nlgn3*^*-/-*^ mice, and *NL3*^*R451C*^ mice (human neuroligin-3 R451C missense mutation) (Hosie et al., 2019; Leembruggen et al., 2020; Margolis et al., 2016). Three out of these four mouse models show changes to the ENS and all mutants demonstrate altered GI function. SERT Ala56 mice have a hypoplastic ENS and SERTKO mice have a hyperplastic ENS (Margolis et al., 2016), both resulting in slower colonic motility. *NL3*^*R451C*^ mice have an increased number of neurons in the SI (Hosie et al., 2019; Leembruggen et al., 2019), while *Nlgn3*^*-/-*^ mice have overall normal numbers of enteric neurons. We find no changes in the number and organization of neurons within the myenteric plexus, but given the function of Caspr2 as a cell-adhesion molecule in neural circuit assembly (Anderson et al., 2012) further investigation is needed to assess whether the connectivity and function, particularly of colonic IPANs, is altered in *Cntnap2*^*-/-*^ mice.

While Cntnap2 was first identified as playing a role in the localization of potassium channels to the juxtaparanodal regions of myelinated axons (Poliak et al., 1999, 2003), considering Cntnap2’s described roles and functions in other systems may shed light on its mechanisms of action in the unmyelinated ENS. Cntnap2 regulates the excitability of DRG sensory neurons by altering Kv1 channel function (Dawes et al., 2018). The absence of Cntnap2 leads to a reduction in overall expression of Kv1.2 channels at the soma membrane of DRG neurons, resulting in altered electrical properties and increased neuronal excitability (Dawes et al., 2018). The gene encoding Kv1.2 (*Kcna2*) is also highly expressed in colonic sensory neurons (Drokhlyansky et al., 2020). Furthermore, similar to the altered cerebellar response to somatosensory stimuli previously reported in *Cntnap2*^*-/-*^ mice (Fernández et al., 2021), IPANs may become hyperexcitable when activated in the absence of Cntnap2, contributing to increased CMC frequency and a shorter time to expulsion during the artificial pellet assay. In contrast, repetitive contractions in isolated empty *Cntnap2* mutant colons were shortened and we attribute this difference in *ex-vivo* colonic motility to the differential activation of sensory neurons (empty colon versus artificial fecal pellet). IPAN-specific manipulations will be instrumental in further investigating the role of Cntnap2 in gut-intrinsic sensory neurons.

By demonstrating altered GI motility, the *Cntnap2*^*-/-*^ mouse model contributes to our understanding of the relationship between ASD and GI dysfunction. Our findings show that, in addition to previously described phenotypes in the CNS and PNS, *Cntnap2*^*-/-*^ mice display changes in colonic motility. Cntnap2’s expression in enteric sensory neurons suggest that sensory dysfunction might contribute to disrupted GI motility. Our findings therefore may have important implications for the diagnosis and treatment of GI symptoms in individuals with ASD.

## 5 Conflict of Interest

The authors declare that the research was conducted in the absence of any commercial or financial relationships that could be construed as a potential conflict of interest.

## 6 Author Contributions

B.G.R. and J.A.K.: conceptualization, data curation, manuscript writing. B.G.R.: data collection, figure preparation, and visualization. B.A.O. and K.M.R.: methodology and analysis. All authors contributed to the article and approved the submitted version.

## 7 Funding

This work was supported by an National Institutes of Health (NIH) NIMH T32 Stanford Neurosciences Program Training Grant, T32MH020016 (B.G.R., K.M.R.); a Bertarelli Foundation Fellowship, the Stanford Master of Science in Medicine Program (B.G.R.); a BP-ENDURE grant from NIH NINDS awarded to the University of Nevada, Reno, R25NS119709 (B.A.O.); the Wu Tsai Neurosciences Institute, the Stanford University Department of Neurosurgery, and research grants from The Shurl and Kay Curci Foundation, and The Firmenich Foundation (J.A.K.).

## 8 Acknowledgments

We thank J. Gomez-Frittelli, R. Hamnett and C. Plant for feedback on the manuscript. We thank V. Lennon (Mayo Clinic) for the HuC/D antibody, J. Sanes (Harvard) for the β-galactosidase antibody, and G. Hennig for the Volumetry software. We thank the Human Pathology/Histology Service Center at Stanford School of Medicine for their slide scanning services. We acknowledge the usage of generative AI technology (ChatGPT May 24 version) for proofreading parts of the discussion section. We have followed all Frontiers guidelines and policies, including verifying the factual accuracy and ensuring plagiarism-free content, and made every reasonable attempt to correct obvious errors.

## 9 Data Availability

Detailed protocols are available upon request. Details of statistical analyses are provided in the Methods section. Datasets are available on request. Correspondence and requests for all materials should be addressed to J.A.K.

## Notes

### Competing Interest Statement

The authors have declared no competing interest.

### Summary of Updates

Now using "Cntnap2" nomenclature instead of “Caspr2.” N-values have been updated in experiments to satisfy power analyses. Higher resolution images have been added to figures. Additional parameters such as CMC Interval have been added.

