## Supplementary Material Figure S1 for "Loss of ASD-Related Molecule Cntnap2 Affects Colonic Motility in Mice"

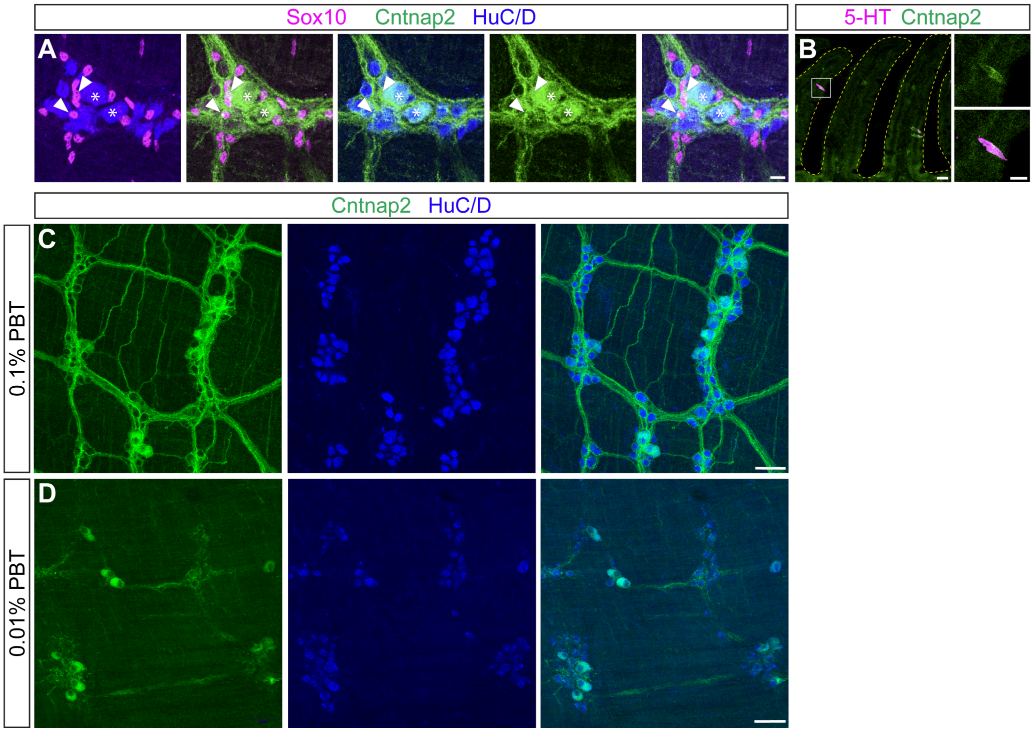


**Figure S1. Cntnap2 colocalizes with a subset of progenitor/glial and enteroendocrine cells. (A)** Cntnap2 (green) colocalizes with a subset of Sox10$^{+}$ cells (magenta) (arrowheads). Enteric neurons labeled with HuC/D (blue). Cntnap2$^{+}$ neurons indicated with asterisks. **(B)** Cntnap2 (green) colocalizes with a subset of 5-HT$^{+}$ cells (magenta) in the epithelium. **(C, D)** Cntnap2$^{+}$ (green) cell bodies and projections are visible with 0.1% PBT (C), while cell body labeling is pronounced with 0.01% PBT (D). Scale bar, (A, B) 10 µm, (C) 50 µm.
